# A green-fluorescent siderophore protects bacterial communities from UV damage

**DOI:** 10.1101/2023.09.26.559457

**Authors:** Özhan Özkaya, Jos Kramer, Tobias Wechsler, Rolf Kümmerli

## Abstract

Sunlight enables virtually all life on earth, but also entails harmful ultraviolet (UV) radiation inducing DNA damage. In response to UV stress, natural selection has favored both curative and preventive measures such as DNA repair mechanisms and UV-absorbing pigments. While UV protection by pigments is well documented in plants, animals and fungi, little is known about their protective role in bacteria. Here, we combine batch-culture and microscopy experiments to show that the siderophore pyoverdine, a fluorescent pigment produced by the opportunistic pathogen *Pseudomonas aeruginosa* offers high-level protection against UV radiation. Our results reveal that bacteria up-regulate pyoverdine production following UV exposure, seemingly as part of a general stress response. We further found that pyoverdine cannot curatively alleviate UV-damage but protects cells preventively and collectively from oncoming UV exposures through its accumulation in the environment. Altogether, our results reveal a new and non-canonical function of this iron-scavenging molecule, demonstrating that pyoverdine acts as a public sunscreen protecting bacterial populations from UV damage. Given that many bacteria produce pigments, such protection might be widespread in species colonizing habitats exposed to UV radiation.

## INTRODUCTION

Sunlight is the energy source that enables virtually all life on earth (Hohmann-Marriott and Blankenship 2011; Rapf and Vaida 2016). However, sunlight also entails UV radiation that causes DNA damage and is thereby harmful for life (Hollósy 2002; Pfeifer et al. 2005). Consequently, many organisms feature adaptations that offer protection against UV radiation or mitigate the damages it causes (Jansen et al. 1998; Hollósy 2002). DNA repair mechanisms are particularly important and occur across all domains of life (Clémenson and Marsolier-Kergoat 2009; Rastogi et al. 2010), while protection against UV radiation typically involves UV-absorbing pigments (Hollósy 2002; Dinkova-Kostova 2008; Solano 2014). Pigment formation is a common prophylactic measure in eucaryotes and occurs in plants (e.g. carotenoids, flavonoids and anthocyanins; Dinkova-Kostova 2008) as well as in animal and fungi (e.g. polyvalent melanins; Butler and Day 1998; Solano 2014). In contrast, much less is known about whether bacteria use pigments as UV protectants, too. Given that they populate a wide range of UV-exposed habitats – such as water surfaces, plant leaves, and host skin (Madigan et al. 2021) – strong selection for UV protection mechanisms could be anticipated in these organisms.

Indeed, many bacterial species produce pigments (Venil et al. 2013). Some of them, like the yellow-brown pigment scytonemin produced by marine cyanobacteria, have been shown to serve as UV protectants (Garcia-Pichel and Castenholz 1991; Sorrels et al. 2009). However, most bacterial pigments are typically examined in the context of functions other than UV protection. For example, staphyloxanthin, the golden pigment produced by *Staphylococcus aureus*, is primarily studied in infections, where it protects cells from oxidative stress imposed by the host immune system (Liu et al. 2005; Clauditz et al. 2006). Violacein, a purple pigment produced by *Chromobacterium violaceum* is often studied in the context of interspecific competition, where it acts as a highly potent anti-bacterial toxin (Choi et al. 2015; Durán et al. 2016). Finally, the group of fluorescent Pseudomonads produces phenazines, blue-green pigments that primarily act as catalyst of redox reactions and toxins in bacterial warfare (Laursen and Nielsen 2004; Mavrodi et al. 2006). Given that bacterial pigments are plentiful and frequently involved in responses to different environmental stressors (Liu and Nizet 2009; Galasso et al. 2017; Pavan et al. 2020; Sajjad et al. 2020), it seems intuitive to ask whether some may also offer protection against UV radiation.

Here, we tackle this question by studying the production of the yellow-green pigment pyoverdine in the opportunistic human pathogen *Pseudomonas aeruginosa*. This versatile bacterium has been isolated from a broad range of habitats, including freshwater, soil and host-associated environments (Crone et al. 2020). Pyoverdine is a siderophore that is secreted into the environment when bioavailable iron is scarce (Visca et al. 2007; Cornelis 2010). Considering its remarkably high affinity for ferric iron and the tight regulation of its synthesis and uptake machinery in response to iron limitation, there is little doubt that the primary function of pyoverdine involves iron acquisition (Visca et al. 2007; Cornelis 2010). However, pyoverdine is excitable by UV radiation and auto-fluorescent, emitting light in the green-yellow spectrum (Meyer 2000). For this reason, it was proposed that pyoverdine together with pyocyanin (a phenazine) could serve as protectants against UV radiation in *P. aeruginosa* (Burke et al. 1990).

In our study, we tested this hypothesis and examined whether UV protection could be a secondary function of this siderophore. First, we tested whether *P. aeruginosa* up-regulates pyoverdine production in response to UV exposure. A relatively strong response under iron-rich conditions (where this pigment is typically not produced) could point towards a regulatory mechanism that controls pyoverdine production outside the context of iron acquisition, but in relation to stressors such as UV radiation. Second, we explored whether the up-regulation of pyoverdine production helps bacteria to mitigate damages incurred during earlier UV exposure or whether it helps to prevent further damage during later UV exposures. In order words, we ask whether the putative upregulation of pyoverdine production offers curative or preventive protection. Third, we tested whether secreted pyoverdine also protects non-producing cells, thereby offering group-level benefits. This scenario can apply when the secreted molecules act preventively as public “sunscreen” protecting not only the producer but also other community members. Finally, we investigated whether individual cells differ in the amount of pyoverdine they make upon UV exposure. One intuitive expectation would be that more damaged cells increase their production of pyoverdine to safeguard themselves against further damage. Alternatively, severely damaged cells with low survival prospects could show a terminal investment (Clutton-Brock 1984; Refardt et al. 2013) by overinvesting in pyoverdine production to altruistically protect less damaged clonemates from future UV exposure.

To address these points, we combined batch-culture experiments with single-cell microscopy and quantified pyoverdine production and its fitness effects under a range of conditions varying in iron availability, the availability of previously produced pyoverdine and UV exposure. Specifically, we measured the effects of all combinations of conditions on the death, survival and growth as well as on pyoverdine production and gene expression of the *P. aeruginosa* PAO1 wildtype strain and its isogenic, pyoverdine-deficient mutant in a full-factorial design.

## RESULTS

### *P. aeruginosa* increases pyoverdine production upon UV exposure regardless of iron limitation

We conducted a series of experiments that all featured the same basic design. We exposed the *P. aeruginosa* wildtype strain PAO1 and its isogenic, pyoverdine-deficient mutant PAO1Δ*pvdD* to three different environmental conditions in a full-factorial design. Specifically, we cultured the bacteria in either iron-limited or iron-rich CAA (casamino acids) medium, with or without supplementation of (previously produced) pyoverdine and with or without prior UV exposure. Replicated bacterial populations were cultured in 96-well plates and incubated in a multi-mode plate reader.

In a first experiment, we measured integrals of population-level pyoverdine production via its natural fluorescence corrected by population growth (Table 1A; Figure 1A; Figure S1). As expected, cultures with the PAO1Δ*pvdD* mutant yielded virtually no fluorescent signal even under iron-limited conditions (0.46 % of wildtype fluorescent levels), confirming the inability of this strain to produce pyoverdine. When focusing on the wildtype PAO1 strain in medium without prior pyoverdine supplementation, we found increased de novo pyoverdine production in UV-exposed compared to non-exposed cultures under both iron-rich and iron-limited conditions (estimate ± SE; iron-limited: 14.24 ± 1.19, t_9.86_ = 11.961, p < 0.001; iron-rich: 4.93 ± 1.19, t_10.08_ = 4.145, p = 0.002). This finding supports our hypothesis that cells respond to UV exposure by producing more pyoverdine. This up-regulation was not observed in medium supplemented with pyoverdine (iron-limited: 2.77 ± 2.51, t_10.01_ = 1.106, p = 0.294; iron-rich: -0.03 ± 0.29, t_10.00_ = -0.101, p = 0.921). These results indicate that cells invest little in de novo production when pyoverdine is already present in the medium (Figure 1A). Altogether, our results show that UV stress triggers increased pyoverdine production regardless of iron availability when the medium itself does not contain previously produced pyoverdine.

**Figure 1.**
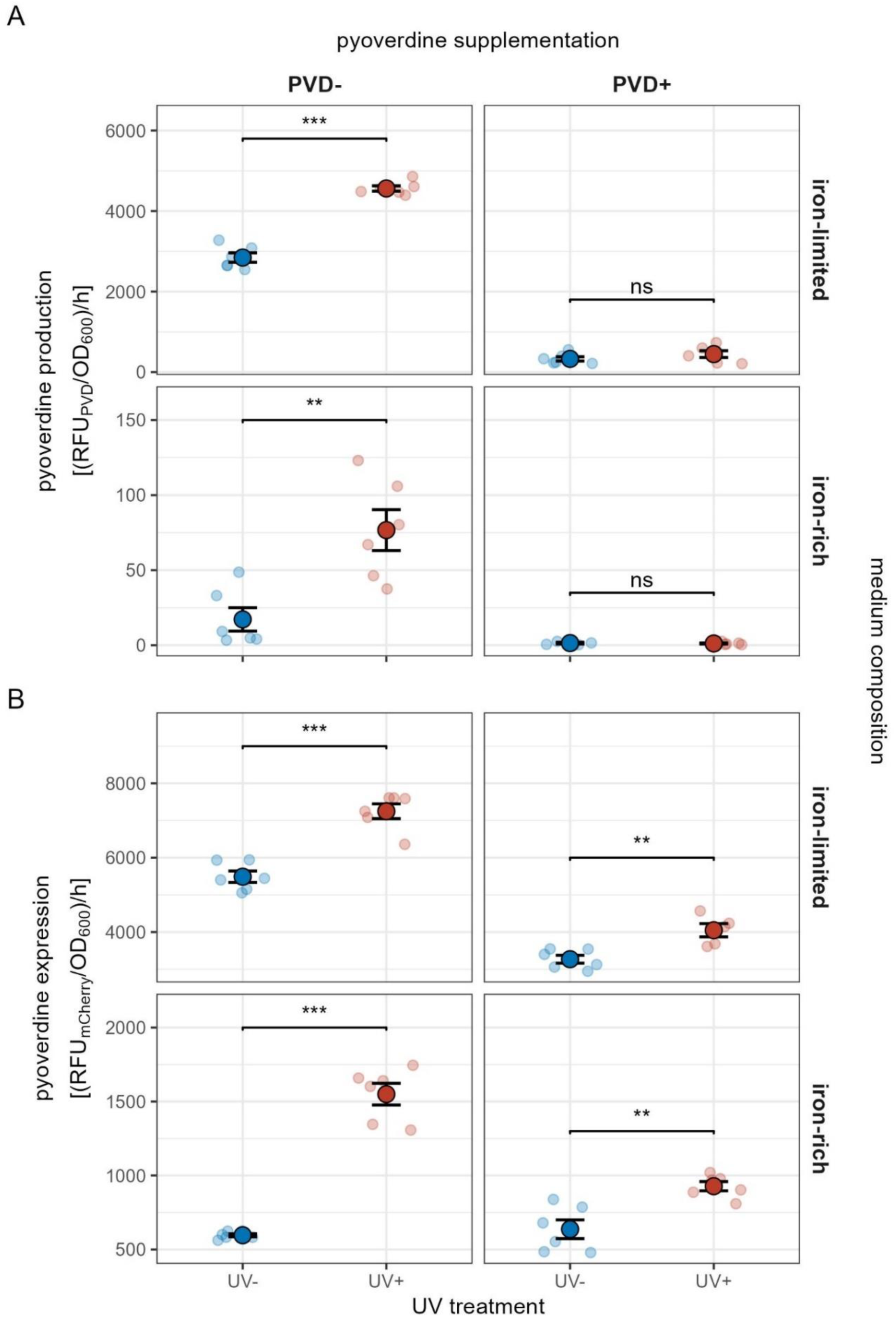
*P. aeruginosa* up-regulates pyoverdine production and synthesis gene expression upon UV exposure regardless of iron availability. (A) Pyoverdine production of PAO1 wildtype monocultures and (B) pyoverdine gene expression of PAO1*pvdA::mCherry* monocultures grown in iron-limited and iron-rich medium with (red) and without (blue) prior UV exposure and prior pyoverdine supplementation. Fluorescence and growth integrals were measured during the exponential phase, as lag phase values were biased by the supplementation of pyoverdine (see Figure S1+S2). The duration of growth phases was obtained by extracting and averaging over the relevant values from Gompertz fits on the data of individual replicates (Figure S3). Pyoverdine production and expression rates were then calculated by first dividing the integral of pyoverdine fluorescence [RFU_PVD_] or mCherry fluorescence [RFU_mCherry_] by the corresponding integral of growth [OD_600_], and then dividing this ratio by the duration of the exponential phase. Small points show individual replicates (n = 5-6), large points and black lines show mean and standard error, respectively. Significance levels of post-hoc comparisons are indicated as follows: * 0.05 ≥ p > 0.01; ** 0.01 ≥ p > 0.001; *** p ≤ 0.001. Note that the y-axis range differs between panels to properly resolve the effect of UV-exposure across all combinations of medium composition and pyoverdine supplementation.

**Table 1.** Effects of treatment conditions on pyoverdine production and fitness. Determinants of (A) the rate of pyoverdine production and (B) the rate of pyoverdine expression in the *P. aeruginosa* PAO1 wildtype as well as (C) growth, and (D) mortality rates of the wildtype and its isogenic, pyoverdine-deficient mutant. Significant p-values are indicated in bold.

| predictor | (A) pyoverdine production rate |  | (B) pyoverdine expression rate |  | (C) growth |  | (D) duration of lag phase |  | (E) mortality rate |  |
| --- | --- | --- | --- | --- | --- | --- | --- | --- | --- | --- |
| | $\chi^2_1$ | p | F<sub>1</sub> | p | $\chi^2_1$ | p | $\chi^2_1$ | p | $\chi^2_1$ | p |
| Supplement (SU) | 380.95 | < 0.001 | 254.14 | < 0.001 | 260.07 | < 0.001 | 58.24 | < 0.001 | 138.29 | < 0.001 |
| Treatment (TR) | 13.33 | < 0.001 | 175.81 | < 0.001 | 136.07 | < 0.001 | 175.05 | < 0.001 | 154.68 | < 0.001 |
| Medium (ME) | 3801.86 | < 0.001 | 3167.30 | < 0.001 | 2078.57 | < 0.001 | 34.98 | < 0.001 | 35.28 | < 0.001 |
| Strain (ST) | - | - | - | - | 4.07 | 0.044 | 5.73 | 0.017 | 1.46 | 0.227 |
| SU : TR | 28.94 | < 0.001 | 24.60 | < 0.001 | 87.59 | < 0.001 | 108.50 | < 0.001 | 7.11 | 0.008 |
| SU : ME | 567.83 | < 0.001 | 111.55 | < 0.001 | 82.66 | < 0.001 | 1.22 | 0.270 | 0.54 | 0.463 |
| TR : ME | 27.20 | < 0.001 | - | - | - | - | 1.80 | 0.180 | 0.02 | 0.902 |
| ST : SU | - | - | - | - | 15.41 | < 0.001 | - | - | - | - |
| ST : ME | - | - | - | - | 33.59 | < 0.001 | - | - | - | - |
| SU : TR : ME | 4.61 | 0.032 | - | - | - | - | 6.92 | 0.009 | 8.15 | 0.004 |
| ST : SU : ME | - | - | - | - | 56.20 | < 0.001 | - | - | - | - |

### *P. aeruginosa* increases PVD gene expression upon UV exposure regardless of iron limitation

While we observed an increase in pyoverdine production upon UV exposure in media without pyoverdine supplementation, the absolute increase was relatively small in iron-rich medium (Figure 1A). However, the fluorescence of pyoverdine is quenched once this molecule binds iron, which suggests that the true extent of up-regulation is masked under iron-rich conditions. To verify that cells indeed show a marked response to UV exposure, we repeated the above experiment using the fluorescent reporter strain PAO1*pvdA::mCherry* to measure the expression of the pyoverdine synthesis gene *pvdA* (Table 1B; Figure 1B; Figure S2).

Our analysis revealed that UV-exposure increases *pvdA* gene expression across all experimental conditions (9.42 ± 0.72, t_41_ = 13.140, p < 0.001). Notably, pyoverdine supplementation generally reduced the magnitude of *pvdA* upregulation, but did so more strongly under iron-limited as opposed to iron-rich conditions (iron-limited: -19.2 ± 1.03, t_41_ = -18.767, p < 0.001; iron-rich: -4.1 ± 1.00, t_41_ = -4.096, p < 0.001; difference: -15.1 ± 1.43, t_41_ = 10.562, p < 0.001). As for pyoverdine fluorescence, *pvdA* expression was lower in iron-rich as compared to iron-limited medium (-40.1 ± 0.72, t_41_ = 56.018, p < 0.001), but the difference in gene expression between these two media (78.6 %) was much less pronounced than the difference in pyoverdine fluorescence (98.3%). Hence, cells do react to UV exposure by markedly increasing their pyoverdine investment even in iron-rich environments where pyoverdine is not needed for iron scavenging (Figure 1B). Altogether, these results indicate that there is a regulatory mechanism induced by stress other than iron limitation that enables cells to ramp up pyoverdine production.

### Pyoverdine buffers the negative growth effects induced by UV exposure

In the next step, we quantified the fitness effects of UV exposure and examined whether pyoverdine helps to alleviate the expected negative impact on (cumulative) growth (Table 1C; Figure 2A; Figure S3). We first focused on growth in media without prior pyoverdine supplementation. Independent of UV exposure, we observed that the pyoverdine mutant grew worse than the wild-type in iron-limited medium (-0.61 ± 0.06, t_20.7_ = -10.840, p < 0.001), while it outgrew the wild-type in iron-rich medium (0.19 ± 0.06, t_11.8_ = 3.009, p = 0.011). These results recover previous findings showing that pyoverdine production is beneficial under iron-limitation, but can come with residual fitness costs under iron-rich conditions (Griffin et al. 2004). More crucially, we found that UV exposure significantly reduced growth in both strains and media (-0.44 ± 0.03, t_50_ = -14.626, p < 0.001). These results show that UV harms the cells, and that the increased pyoverdine production observed in the wildtype strain upon UV exposure (Figure 1) does not seem to alleviate that harm. In both strains and media, we observed that the supplemented pyoverdine had an overall positive effect on growth (0.59 ± 0.03, t_53_ = 19.890, p < 0.001) as it accelerates iron acquisition. But crucially, the growth increase was much more pronounced after UV exposure (0.56 ± 0.06, t_50_ = 9.359, p < 0.001), resulting in an almost complete rescue of growth in both strains regardless of iron availability (Figure 2A).

**Figure 2.**
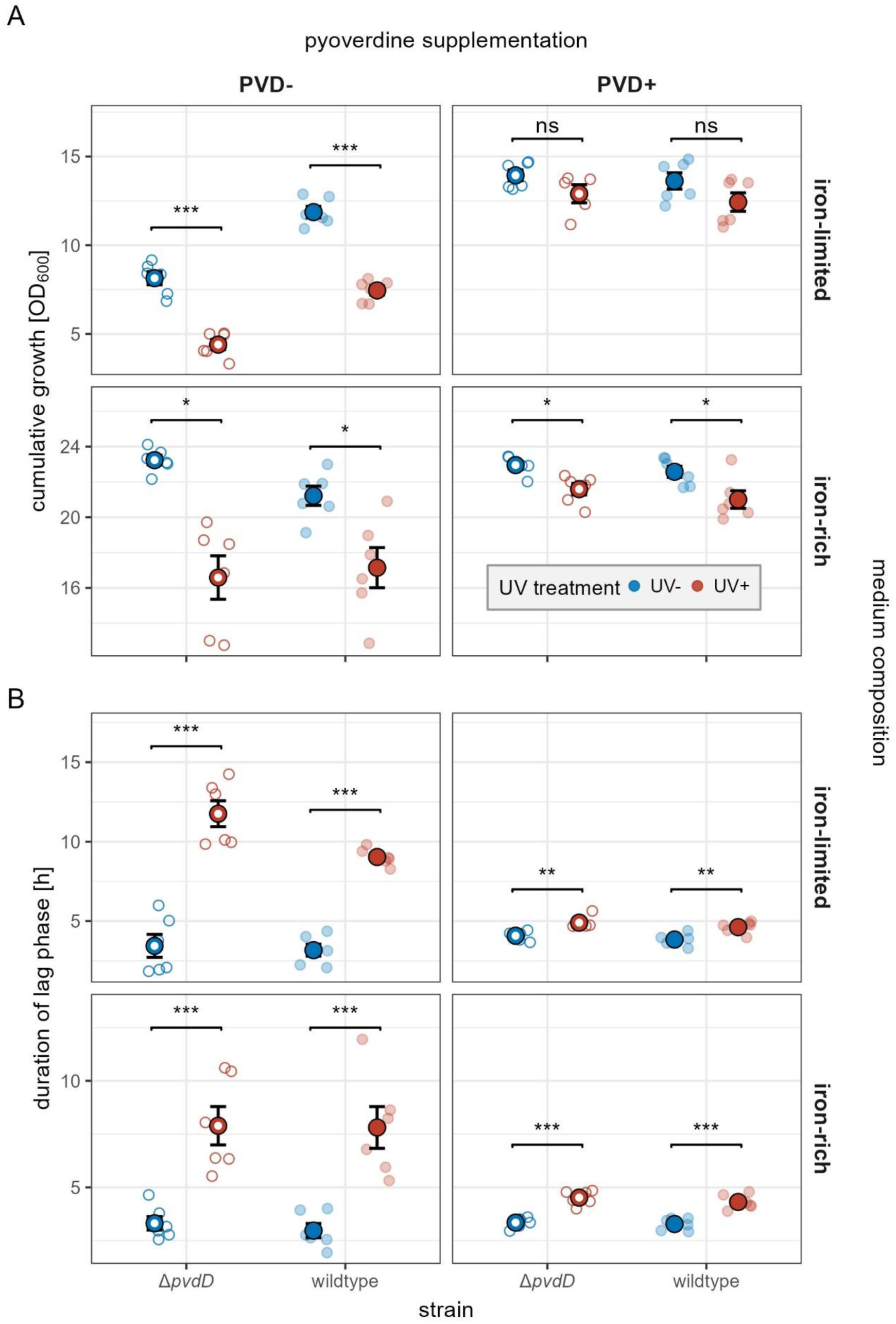
Pyoverdine supplementation prior to UV exposure rescues bacterial population growth and shortens the lag-phase regardless of iron availability and strain identity. (A) Cumulative growth and (B) lag-phase duration of PAO1 wildtype (solid circle) and PAO1 Δ*pvdD* (open circle) monocultures in iron-limited and iron-rich medium with (red) or without (blue) prior UV exposure and with or without prior pyoverdine supplementation. Growth integrals were measured over the entire duration of the experiment (24h). Lag phase durations were obtained from Gompertz fits on the data of individual replicates (Figure S3). Small circles are individual replicates (n = 5-6), large circles and black lines show mean and standard error. Significance levels of post-hoc comparisons are indicated as follows: * 0.05 ≥ p > 0.01; ** 0.01 ≥ p > 0.001; *** p ≤ 0.001. Note that the y-axis range differs between panels to properly resolve variation between strains across treatment combinations.

To investigate the impact of UV exposure on growth dynamics, we analysed its effects on the duration of the lag phase (Table 1D; Figure 2B; Figure S3). We found that UV exposure substantially increased the lag phase in both strains and under all conditions (0.73 ± 0.05, t_49.4_ = 15.421, p < 0.001). In line with a protective function of pyoverdine, we observed that pyoverdine supplementation decreased the lag phase after UV exposure (-0.86 ± 0.07, t_49.5_ = -12.769, p < 0.001). By contrast, pyoverdine supplementation in the absence of UV stress had no effect on the duration of the lag phase under iron-rich conditions (0.06 ± 0.10, t_49.3_ = 0.624, p = 0.536), while it slightly prolonged the lag phase under iron-limited conditions (0.20 ± 0.10, t_49.3_ = 2.147, p = 0.037). Independent of all these effects, the pyoverdine mutant generally featured a longer lag-phase than the wildtype (0.05 ± 0.02, t_47.3_ = 2.394, p = 0.021). Altogether, these findings demonstrate that (i) pyoverdine needs to be present prior to UV exposure to exert an immediate protective function, and (ii) that this preventive protection also accrues to pyoverdine non-producers.

### Pyoverdine reduces cell death caused by UV exposure

Above, we examined the effect of UV exposure on population growth parameters. While they are useful proxies for fitness, they do not inform us on whether UV exposure kills cells or just reduces their division rate. To explore the effects of UV radiation on cell mortality, we developed a dynamic live-dead (propidium iodide) staining assay that allowed us to monitor relative cell death over time in batch cultures (Table 1E; Figure 3). Under all culturing conditions, we observed a relatively high prevalence of dead cells at the beginning of the experiment for both the wildtype and the pyoverdine non-producer (Figure S4). This is expected as we started our experiments with cells from stationary overnight cultures, which typically contain dead cells.

**Figure 3.**
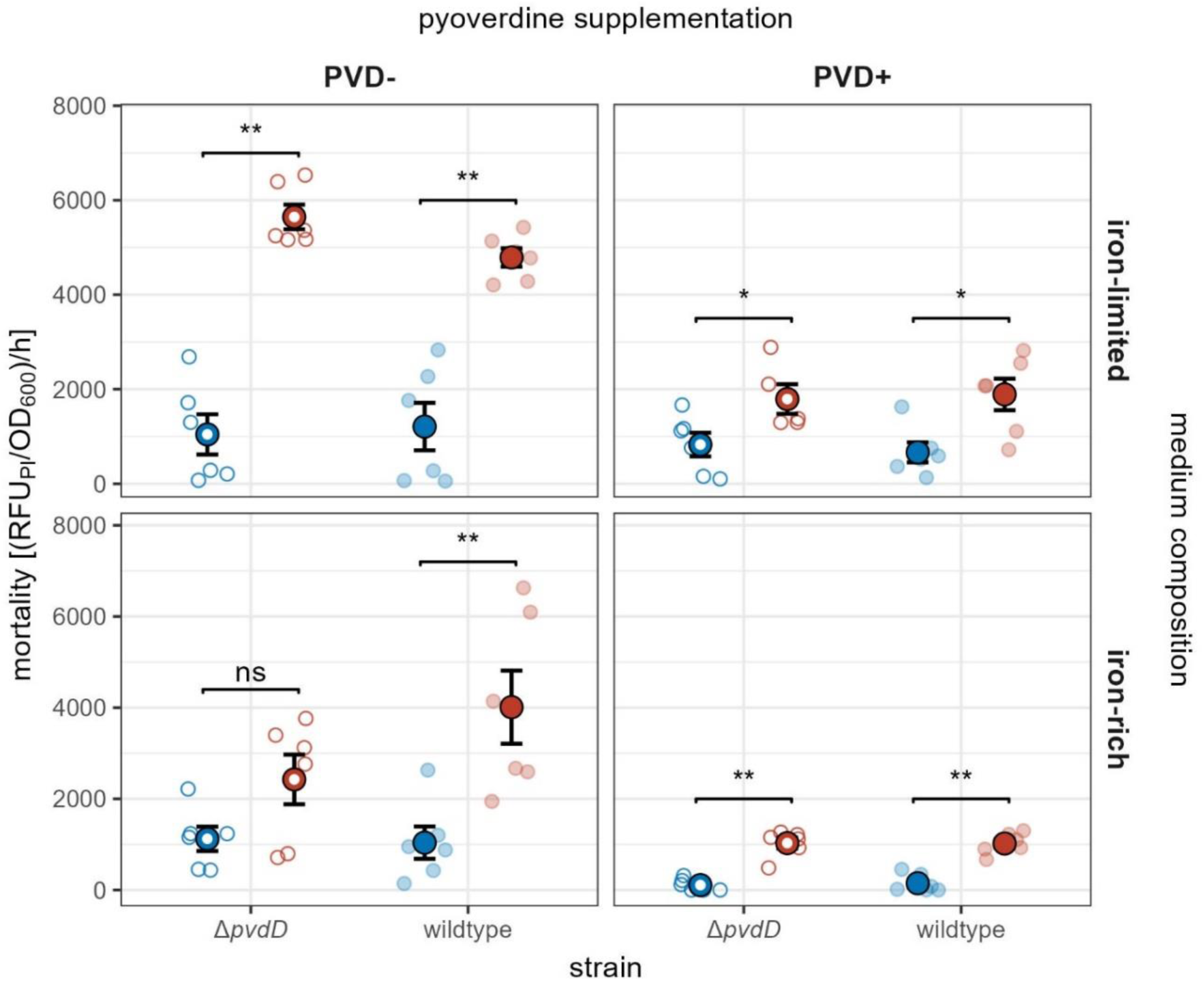
Pyoverdine supplementation prior to UV exposure prevents UV-related cell death. Mortality determined by propidium iodide (PI) dead-cell staining of PAO1 wildtype (solid circle) and PAO1 Δ*pvdD* (open circle) monocultures growing in iron-limited and iron-rich medium with (red) or without (blue) prior UV exposure and pyoverdine supplementation. Fluorescence and growth integrals were measured during the lag phase (Figure S3), during which the impact of UV exposure on cell death is expected to be high. Lag phase durations were obtained by extracting and averaging over the relevant values from Gompertz fits on the data of individual replicates. Mortality was then quantified by first dividing the integral of PI fluorescence [RFU_PI_] by the integral of growth [OD_60_0], and then dividing this ratio by the duration of the lag phase. Small circles are individual replicates (n=5-6), large circles and black lines show mean and standard error. Significance levels of post-hoc comparisons are indicated as follows: * 0.05 ≥ p > 0.01; ** 0.01 ≥ p > 0.001; *** p ≤ 0.001.

In the absence of UV exposure, the prevalence of dead cells quickly fell to the detection limit before slowly starting to rise again as cultures approached stationary phase. By contrast, dead cells occurred at high abundance over many hours of the experiment in cultures exposed to UV (Figure S4). When focusing on the initial growth (i.e. the lag) phase, during which the impact of UV exposure on cell death is expected to be highest, these patterns translated into substantially increased mortality rates after UV exposure in media without pyoverdine supplementation (iron-limited: 43.4 ± 5.45, t_11.6_= 7.962, p <0.001; iron-rich: 23.4 ± 5.78, t_19.5_ = 4.051, p < 0.001). Crucially, when the media had been supplemented with pyoverdine before UV exposure, we observed greatly reduced killing (iron-limited: -29.74 ± 3.01, t_12.9_ = -9.882, p <0.001; iron-rich: -22.71 ± 4.92, t_12.4_ = -4.613, p < 0.001). Note that all these patterns applied equally to both the mutant and the wildtype strains. Overall, these results demonstrate that UV exposure kills a high fraction of cells in the population – and that this killing is almost entirely prevented by pyoverdine supplemented prior to UV exposure (Figure 3).

### Heavily damaged cells overproduce pyoverdine upon UV exposure under iron-rich conditions

Until now, we have shown that previously secreted pyoverdine in the medium protects bacterial cells from UV radiation. However, we still do not understand why bacteria up-regulate pyoverdine production upon UV exposure as this up-regulation does not seem to offer immediate protection to the producers. From an inclusive fitness perspective (West et al. 2006), two (non-mutually exclusive) explanations are conceivable. First, UV-damaged cells might up-regulate pyoverdine production for their own benefit, for instance to minimize the likelihood of sustaining even more damage during future UV exposures. Second, damaged cells that are likely to die might cooperatively up-regulate pyoverdine production to increase the concentration of pyoverdine in the environment. By doing so, they would not benefit themselves but increase future protection levels for their (less damaged) clonemates (Lehmann 2007). In both scenarios, pyoverdine production is expected to increase with the severity of UV-induced damage.

To test this prediction, we used time-lapse microscopy to follow the fate of individual cell lineages (i.e., founder cells and their progeny) growing on agarose patches. We observed four distinct phenotypes that can plausibly be linked to the degree of UV-induced damage (Figure 4A, Video S1): [A] healthy cells that started to divide, leading to the formation of microcolonies; [B] moderately damaged cells that did not divide, but remained intact over the duration of the experiment; [C] severely damaged cells that started to divide, but eventually underwent explosive cell lysis (Turnbull et al. 2016), leading to the death of all cells in the microcolony; and [D] critically damaged cells that did not divide and underwent cell lysis. When cells were not exposed to UV, the healthy phenotype [A] predominated (constituting 71% of cell lineages). When exposing cells to UV, the damaged phenotypes [B], [C] and [D] rose to high frequency (constituting 89% of cell lineages; difference between UV treatments, chi-square test: χ^2^ = 247.8, p < 0.001; Figure S5). These results demonstrate the substantial fitness costs of UV exposure both in terms of cell death (lysis) and growth arrest.

**Figure 4.**
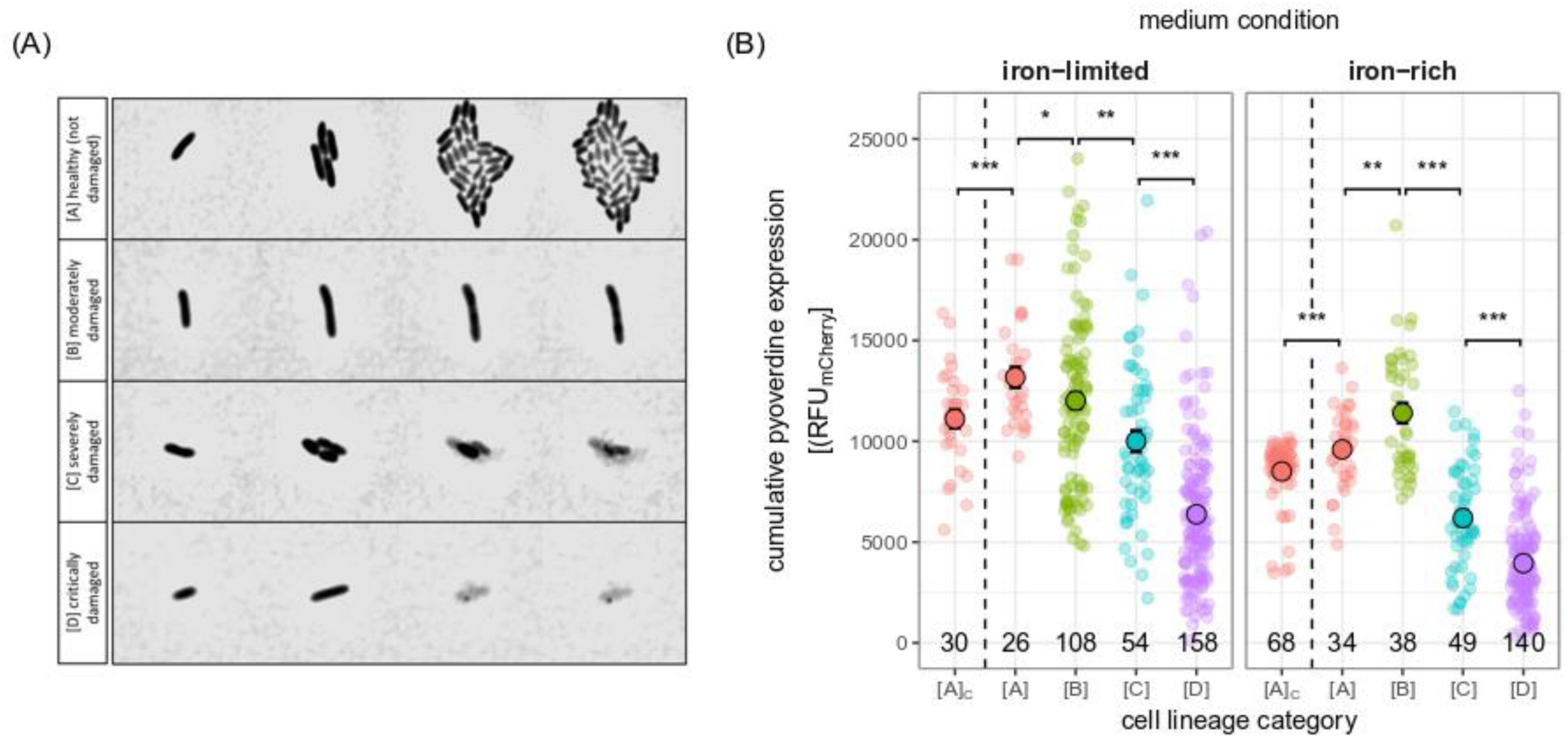
Cells with more severe UV damage express pyoverdine at lower levels. (A) Different cell lineage types varying in the degree of apparent UV-induced damage: [A] healthy cells start to divide and form microcolonies; [B] moderately damaged cells do not divide, but remain intact over the duration of the experiment; [C] severely damaged cells start to divide, but eventually undergo explosive cell lysis, leading to the death of all cells in the microcolony; [D] critically damaged cells do not divide and eventually undergo explosive cell lysis. (B) Cumulative pyoverdine expression of UV-exposed cell lineages that vary in the degree of apparent UV damage, measured under iron-limited and iron-rich conditions. Pyoverdine expression was quantified as the integral of mCherry fluorescence over the duration of the experiment (8h). Small circles are individual replicates, and either represent the signal of individual cells (for non-dividing types [B] and [D] featuring moderate or critical damage) or the average signal of all cells belonging to the same lineage (for dividing types [A] and [C] featuring no or severe damage). [A]C are healthy cells from cultures without UV exposure and serve as a negative control. Large circles and black lines show mean and standard error. Numbers within panels indicate sample sizes. Significance levels of post-hoc comparisons are indicated as follows: * 0.05 ≥ p > 0.01; ** 0.01 ≥ p > 0.001; *** p ≤ 0.001.

We then compared the cumulative cell lineage expression of the *pvdA* synthesis gene. In the absence of UV exposure, we observed an expectably lower *pvdA* expression of healthy [A] lineages in iron-rich compared to iron-poor medium (t-test: t_39.4_ = -5.32, p < 0.001; Figure S5). Next, we compared *pvdA* expression of healthy [A] lineages between UV-treatments and found that expression levels were increased after UV exposure independent of iron limitation (7.17 ± 1.74, t_155_ = 4.116, p < 0.001; interaction: F_1_ = 1.120, p = 0.292; Figure 4B). To test the hypothesis that more damaged cells over-invest in pyoverdine production, we compared *pvdA* expression in healthy [A] versus damaged cell lineages after UV exposure (Table S1A; Figure 4B). Under iron limitation, we found no support for our hypothesis as healthy [A] lineages expressed more pyoverdine than the damaged types [B], [C] and [D] (Table S2). Under iron-rich conditions, pyoverdine expression peaked for [B] lineages, which had both higher expression levels than the more damaged types [C] and [D] and the healthy phenotype [A] (Table S2). Note that the above analyses focus on the cumulative expression of pyoverdine, such that the expression levels of the heavily damaged (lysing) [C] and [D] lineages might be underestimated as they die before the end of the experiment. Consequently, we repeated our analyses focusing on rates of pyoverdine expression. These analyses revealed that the heavily damaged [C] and [D] types expressed pyoverdine at the same or lower rates than the healthier [B] and [A] lineages during their lifetime (see the Supplementary Analysis; Table S1B; Figure S6). Taken together, our results provide no support for the hypothesis that the heavily damaged cell types [C] and [D] over-invest in pyoverdine to altruistically protect their less damaged clonemates from future UV damage. In contrast, moderately damaged cells [B] showed such a pattern of over-investment under iron-rich conditions. Given that these cells survive, the observed response might result in a combination of self-protection and increased protection of clonemates.

## DISCUSSION

Although bacteria populate a wide range of UV-exposed habitats, ranging from water surfaces to plant leaves and host skin, the question of how they protect themselves from UV damage has received scant attention. Here, we tested whether pyoverdine (a siderophore pigment) produced by the opportunistic human pathogen *P. aeruginosa* can exert a curative or offer a preventive function to protect producers and other cells in the vicinity from UV radiation. We found that bacterial populations upregulated both pyoverdine gene expression and production in response to UV exposure. However, this response had no curative effect as bacterial fitness was similarly compromised by UV exposure regardless of whether bacteria were able to produce pyoverdine or not. Conversely, previously secreted (and experimentally supplemented) pyoverdine offered an almost complete preventive protection from UV radiation, substantially reducing post-exposure mortality in both producers and non-producers. At the single-cell level, we found that pyoverdine expression upon UV exposure is heterogeneous. In particular, expression peaked in moderately damaged cells but was reduced in heavily damaged cells that eventually underwent lysis. These results reveal that the upregulation of pyoverdine as a preventive protection measure is primarily triggered in healthy and moderately damaged cells that have future fitness prospects. Altogether, our findings demonstrate that pyoverdine is regulated in response to UV exposure and can act as a public sunscreen protecting bacterial populations from UV damage.

Our findings show that the benefits of pyoverdine production under UV stress do not arise as a mere by-product of investment into the iron-scavenging function of this siderophore because pyoverdine expression is upregulated after UV exposure even under iron-rich conditions. In the context of its canonical, iron-acquisition function, pyoverdine production is up-regulated when bioavailable iron is scarce (Kümmerli et al. 2009). The secreted molecules act as chelators scavenging otherwise inaccessible (insoluble or host-bound) iron for the metabolism of the producer and other cells with matching uptake receptors (Cornelis 2010; Kramer et al. 2020). Here, we show that pyoverdine has a non-canonical function involving the protection of cells from damage caused by UV stress. Our findings extend the list of pyoverdine’s non-canonical functions that also include toxic metal sequestration and intracellular/cell-to-cell signaling (Lamont et al. 2002; Braud et al. 2010).

At the mechanistic level, our results suggest that pyoverdine protects cells from UV light by reflecting/absorbing UV radiation before it reaches the cells (preventive protection), and not by mitigating the damage induced by UV (curative protection). Our results contrast with a recent study by Jin and colleagues (Jin et al. 2018) who showed curative effects of pyoverdine in response to oxidative stress. In their study, pyoverdine retained in the periplasm protected cells from oxidative damage by preventing the formation of reactive oxygen species (ROS) following exposure to visible light or antibiotics. Although UV radiation can also promote ROS formation (Rastogi et al. 2010), our population-level data provide no support for such a curative role of pyoverdine following UV exposure. Instead, our data indicate that pyoverdine exerts a preventive function and thus needs to be present in the environment prior to UV exposure to reduce negative effects on survival and growth. We note, however, that the preventive ‘sunscreen’ protection and the curative protection against ROS formation are non-mutually exclusive and potentially complementary mechanisms that likely come with different costs and benefits depending on prior levels of secreted pyoverdine and the nature of the environmental stress. On a more general level, these dual benefits suggests that the upregulation of pyoverdine might be part of a general stress response, a hypothesis that is also supported by the observation of increased pyoverdine expression after exposure of *P. aeruginosa* to competitor supernatants (Leinweber et al. 2023). This hypothesis requires careful testing by investigating how pyoverdine expression and transport are fine-tuned in response to different environmental stressors.

Our findings demonstrate that pyoverdine can act as a secreted ‘sunscreen’ protecting producers and other community members from UV-induced damage. Population-level benefits of secreted compounds are often substantial and range from the inactivation of antibiotics through the secretion of β-lactamases (Domingues et al. 2017; Bush 2018) to the detoxification of heavy-metal contaminated environments through the secretion of siderophores (Braud et al. 2010; Hesse et al. 2018). However, it often remains unclear whether producers secrete these ‘public goods’ to primarily reap direct benefits for themselves, or whether secretion is an altruistic behavior that indirectly benefits the producer by increasing the fitness of clonemates. Using single-cell microscopy, we show that cellular variation in UV-induced pyoverdine expression is likely not a consequence of altruistic terminal investment, but rather reflects a reduced capability of more heavily damaged cells to ramp up pyoverdine expression after UV exposure. In particular, we found that heavily damaged cell lineages, which eventually underwent cell lysis, showed the lowest levels of pyoverdine expression. This indicates that beyond a certain level of damage, cells no longer over-invest in pyoverdine production, neither for their own nor for the altruistic protection of clonemates. By contrast, cells experiencing no apparent, or moderate damage (non-lysing cells) showed increased pyoverdine expression upon UV exposure. The response of these cells could involve both direct (increased self-protection from future UV exposures) and indirect fitness benefits by protecting their kin.

Although we focused on *P. aeruginosa*, our results on the role of pyoverdine in UV protection are likely of broader relevance. This is because the chromophore of pyoverdine is highly conserved, suggesting that its UV-protective function might apply to all fluorescent pseudomonads, which express pyoverdine as their primary siderophore (Cornelis 2010). Pyoverdine molecules are very durable (Kümmerli and Brown 2010) and likely accumulate to concentrations sufficient for sunscreen protection in at least some natural settings. If such UV protection is relevant for fitness, we predict that pyoverdine production should vary across habitats as a function of UV exposure. Specifically, we would expect *Pseudomonas* species living in soil to produce less pyoverdine than species living on plant leaves or in surface water, where exposure to UV radiation is high. In support of this prediction, we previously showed that pseudomonads isolated from freshwater (pond) samples produce significantly more pyoverdine that isolates from soil samples (Butaitė et al. 2018). However, these environments differ in many aspects other than UV-exposure that could also affect pyoverdine production. Confirmation of a general role of pyoverdine-mediated sunscreen protection therefore requires experiments that quantify pyoverdine concentrations in natural settings and demonstrate their fitness benefits upon exposure to naturally occurring intensities of UV light in different habitats.

In conclusion, we demonstrated a novel non-canonical function of the siderophore pyoverdine, acting as a public sunscreen protecting communities of *P. aeruginosa* bacteria from UV damage. Our results have three general implications: From an adaptationist perspective, they first show that a single molecule can fulfill multiple functions both by performing different tasks (e.g., chelating iron vs. absorbing and reflecting UV light) as well as by performing the same task in different contexts (e.g. chelating iron for metabolism under iron-limitation vs. chelating iron to prevent ROS formation under oxidative stress). Second, they highlight that the benefits of the pyoverdine-mediated sunscreen protection are public, suggesting that species with limited means of UV protection might be able to survive in some environments by exploiting the public UV protection measures of other community members. Finally, our results also have implications for virulence evolution. Specifically, pyoverdine is a virulence factor in infections (Granato et al. 2016), and the question is how such virulence traits can be maintained outside the host (Brown et al. 2012). Our results provide an answer by showing that molecules [e.g., pyoverdine] contributing to virulence during infections can fulfill additional functions in natural environments [e.g., UV protection]. This multifunctionality can both prevent evolutionary deterioration of the virulence trait outside the host and allow opportunistic pathogens to flexibly colonize and thrive in a variety of habitats.

## MATERIALS & METHODS

### Strains, pre-culture conditions, and media

We used three strains of *Pseudomonas aeruginosa* (ATCC 15692): (i) PAO1 WT, our focal wild-type strain that produces pyoverdine as its primary siderophore; (ii) PAO1Δ*pvdD*, an isogenic mutant of our wild-type strain that is unable to produce pyoverdine and serves as a negative control; and (iii) PAO1*pvdA::mCherry*, a reporter strain carrying a promotor-*mCherry* fusion (chromosomal insertion; attTn7::Ptac-mCherry) that allows quantification of the expression of the pyoverdine synthesis gene *pvdA* based on the fluorescence of the marker protein mCherry (details in Rezzoagli et al. 2019). We always grew pre-cultures of these strains in 50 ml Falcon tubes containing 8 ml of Lysogeny Broth (LB) at 37°C under shaking conditions (170 rpm) for 24 hours. Prior to their use in our experiments, we washed pre-cultures three times in 0.85% NaCl solution, and adjusted them to the same optical density (OD_600_= 1; measured at 600 nm using an Infinite M200 Pro microplate reader [Tecan Trading AG, Switzerland]). In our experiments, we grew the strains in (i) iron-limited CAA medium (5 g casamino acids, 1.18 g K_2_HPO_4_·3H_2_O and 0.25 g MgSO_4_·7H_2_O per litre, supplemented with 25 mM HEPES buffer and 200 μM 2,2’-bipyridine, a strong iron chelator), and (ii) an iron-rich control medium, also consisting of CAA but supplemented with 40 μM FeCl_3_.

### Batch-culture experiments

We performed two experiments to investigate the effects of UV radiation on pyoverdine production and population growth. In the first experiment, we quantified the total biomass, pyoverdine production, and ratio of dead over live cells of both the PAO1 WT and its isogenic PAO1Δ*pvdD* mutant after growth in either iron-rich or iron-limited medium, with or without pyoverdine supplementation, and with or without prior UV exposure. To collect the pyoverdine required for supplementation, we grew the PAO1 wild-type in 8 ml iron-limited medium for 24h as described above, and then harvested the (pyoverdine-containing) supernatant by first pelleting the cells (via centrifugation at 170000 rpm for 6 min) and then filter-sterilizing the liquid phase three times (using a 0.22 µm filter). We then mixed the resulting sterile supernatant with iron-rich or iron-limited medium (at a ratio of 1:5) to create master mixes of our pyoverdine-supplemented media. To obtain master mixes of unsupplemented media, we mixed iron-limited or iron-rich medium with sterile 0.85% NaCl solution. To be able to differentiate between dead and live cells, we added propidium iodide (PI) at a final concentration of 5 µg/mL to each of the master mixes. To initiate the experiment, we inoculated washed and OD-adjusted cultures of the PAO1 WT and mutant strains into the four different master mixes in a 96-well plate (2 µl culture and 200 µl medium per well). We inoculated three independent replicates per plate and set up two plates on different days, resulting in six biological replicates per condition. Immediately after inoculation, we exposed one half of the plate comprising replicates of all relevant treatment combinations to UV light (Philips TUV TL-D 30W G13; emission peak at 253.7 nm) for 20 seconds under sterile conditions (the other half of the plate, which comprised all treatment combinations without UV exposure, was covered with aluminum foil to prevent exposure). Finally, we incubated the 96-well plate at 37°C in the plate reader, and quantified population growth (as OD_600_), pyoverdine fluorescence (as relative fluorescence units RFU_PVD_, excitation|emission at 400|460 nm), and the amount of dead cells (as relative fluorescent units RFU_PI_, excitation|emission at 493|646 nm) every 15.8 min (after 120 seconds of rigorous shaking) for the next 24 hours.

In the second experiment, we quantified growth and pyoverdine expression in our reporter strain. This experiment featured the same setup as the previous experiment, with two exceptions: (i) we used PAO1*pvdA::mCherry* instead of PAO1 WT to be able to measure pyoverdine transcription, and (ii) we omitted propidium iodide life-dead stain from all master mixes. This latter change was necessary, because the fluorescent signal of propidium iodide overlaps with the fluorescence of the mCherry marker (which is measured as RFU_mCherry_, excitation|emission at 588|620 nm). For both experiments, we estimated growth curve parameters (Figure S3) using a modified Gompertz equation (Zwietering et al. 1990). We then calculated integrals of pyoverdine production and expression as well as growth integrals and integrals of mortality over the relevant growth phases. For the analysis of pyoverdine production and expression, we calculated integrals over the exponential phase, as lag phase values were biased by the supplementation of pyoverdine in some treatment combinations (Figure S1+S2). For the analysis of growth, we calculated integrals over the whole duration of the experiment. Mortality rates are expected to be highest directly after UV exposure, and we therefore calculated integrals of mortality rate over the duration of the lag phase only. Because growth phase durations differed between wildtype and mutant (see the Results and Figure S3), we divided integrals by the time over which they were measured before using them in our statistical analyses.

### Microscopy experiments

We used time-lapse microscopy to study heterogeneity in pyoverdine production and in the expression of the *pvdA* synthesis gene in our reporter strain after UV exposure. To this end, we first grew pre-cultures of the PAO1*pvdA::mCherry* reporter strain, and then inoculated them on agarose patches on microscopy slides with a custom-made design. Specifically, we adopted a microscopy slide setting based on a previous study (Weigert and Kümmerli 2017) and modified it to accommodate two separate Gene frames (Thermo Fisher Scientific) on the same microscopy slide (Figure S7). A Gene frame is a double-sided adhesive material that creates an air-sealed rectangular space for agarose patches between the microscopy slide and the cover glass. Using two gene frames on the same microscopy slide allowed us to test two different media on the same microscopy slide. Per Gene frame, we used 200 µl of iron-limited or iron-rich medium (see above), solidified with 1% (mg/ml) agarose. Within each Gene frame, we created four separate square agarose patches on each corner by cutting out a cross-shaped corridor between them, thereby ensuring both the isolation of cells among different agarose patches and sufficient oxygen flow to allow cell survival and growth (Figure S7).

To start the experiment, we inoculated the two top and the two bottom patches in each gene frame with 2 µl of washed and adjusted culture of PAO1*pvdA::mCherry* and PAO1Δ*pvdD* (which was only used as an internal control), respectively. In analogy to the batch culture experiments, we covered one half of the microscopy slide (the first Gene frame) with aluminium foil while exposing the other half (the second Gene frame) to UV light for three seconds. Note that we used a shorter duration of UV exposure than in the batch culture experiments, as cells were not covered by liquid medium, and were thus much more vulnerable to UV exposure on the agarose patches. Directly after UV exposure, we sealed the Gene frames airtight with coverslips, and then used a Widefield – Olympus ScanR HCS microscope (Center for Microscopy and Image Analysis, University of Zurich) to detect and follow cells and micro-colonies using phase contrast imaging. We further measured *pvdA* gene expression through mCherry fluorescence (via the TRITC filter; excitation|emission: 550 nm|580nm) and pyoverdine fluorescence visible in the periplasm (via the DAPI filter; excitation|emission: 359 nm|457nm). The two measures are highly correlated under iron-limited conditions (5.878e^-02^ ± 6.242e^-03^, t_28_ = 9.416, p < 0.001, R^2^ = 0.75). For the main analysis, we used the *pvdA* gene expression data because it reliably measures cellular pyoverdine investment levels while pyoverdine auto-fluorescence in the periplasm is the sum of self-produced and imported pyoverdine. Images were taken at two different fields of view (each corresponding to a technical replicate) on each agarose patch every 10 minutes for 11 hours. Throughout the time-lapse measurements, the slides were incubated at 37°C in the microscope’s incubation chamber.

For image analysis, we developed a pipeline based on the open-source segmentation and tracking softwares SuperSegger (Stylianidou et al. 2016) and ImageJ (https://imagej.net/; Schneider et al. 2012). We first imported all time-lapse recordings into ImageJ for manual review and then exported individual images (one Image per time-point and channel) for further analysis to SuperSegger. We only used the first eight hours of each experiment, as most micro-colonies merged, and the fields of view became overcrowded after that timepoint. We used SuperSegger to (i) segment and identify each single cell based on the phase contrast images, (ii) track each cell in space and time, (iii) register each cell division event to reconstruct cell lineage trees, and (iv) quantify *pvdA* gene expression as well as pyoverdine content in the periplasm of each single cell. Cell lineage information and mean fluorescence values for each individual cell were then transferred to R for further analysis. We grouped cells into four life history categories based on the occurrence of cell division and explosive cell lysis (Video S1 and Figure 4A): [A] dividing, non-lysing cells (present in all frames); [B] non-dividing, non-lysing cells (present in all frames); [C] dividing and lysing cells (present in the first frame, lineage goes extinct in subsequent frames); [D] non-dividing, lysing cells (present in the first frame, lyses in a subsequent frame). We used two different approaches to calculate *pvdA* expression per cell and cell lineage. First, we measured the integral of the mCherry signal across all frames, yielding a cumulative measure of gene expression. Second, we calculated the integral of the mCherry signal divided by the number of frames in which the signal was detected, returning a rate of pyoverdine gene expression. Note that both measures are based on the mCherry signal of either a single cell (categories [B] and [D]), or the average of all cells belonging to the same lineage [categories [A] and [C]).

### Statistical Analyses

We used two linear models (LMs) and five generalized least square (GLS) models for statistical analyses in R 4.2.0 (www.r-project.org). One GLS model and one LM were fitted using, respectively, the per-capita rate of pyoverdine production or pyoverdine expression of the PAO1 wildtype as response variable, and UV-treatment (UV+ or UV-), pyoverdine supplementation (PVD+ or PVD-) as well as medium (iron-limited or iron-rich) as explanatory factors. To analyze the effects of UV exposure on growth and cell death, we fitted three additional GLS models using, respectively, (cumulative) growth, the duration of the lag phase, and mortality rate as response variable, and UV treatment, pyoverdine supplementation, medium and strain (wild type vs. pyoverdine mutant) as explanatory factors. Finally, we analyzed the single-cell data in a LM and an additional GLS model. The LM was fitted on data of healthy cell lineages and tested whether their cumulative pyoverdine expression depended on UV treatment or medium. The GLS model focused on data from UV-exposed cells and tested whether the cumulative pyoverdine expression depended on the medium or the cell lineage type (categorical; healthy [A], moderately damaged [B], severely damaged [C], or critically damaged [D]). We fitted GLS models instead of LMs whenever the inspection of the initially fit LMs (using functions implemented in the performance package; Lüdecke et al. 2021) indicated a deviation from residual homogeneity. In the GLS models, we accounted for this variance heterogeneity by selecting an appropriate variance structure (implemented via the weights-argument of the ‘gls’ function of the nlme package; Pinheiro et al. 2023) based on model comparisons via AIC (Zuur et al. 2009). The p-values of effects in all models were obtained using the ‘Anova’ function (car package; Fox & Weisberg 2019). We used the emmeans package (Lenth 2021) to perform post-hoc analyses and adjusted p-values for multiple testing (n_test_>2) using the false discovery rate. All models were initially fitted with all interaction terms. Each model was then simplified stepwise by dropping non-significant interaction terms (p > 0.05). The structure of all final models is detailed in Table S4. Finally, note that we excluded two replicates that grew to much higher density than the other replicates in the same treatment (>1.5-fold increased density) from all analyses, as their inclusion caused problems in our statistical models.

## AUTHOR CONTRIBUTION STATEMENT

Conceptualization and design: ÖÖ and RK; lab work: ÖÖ; bioinformatics: TW and JK; statistical analyses: JK; interpretation of the results: ÖÖ, JK, and RK; writing (first draft): ÖÖ, JK and RK; writing (elaboration): JK and RK.

## ACKNOWLEDGEMENTS

We thank Amar Lekovic for helping with the batch culture parts of this study. This research was supported by the European Research Council (ERC) under the European Union’s Horizon 2020 research and innovation program (grant agreement no. 681295) and the Swiss National Science Foundation (31003A_182499) to RK as well as by a UZH Postdoc grant (Forschungskredit; K-74822-01) and the University Research Priority Program “Evolution in Action” to ÖÖ.

