## Supplementary Material for "A green-fluorescent siderophore protects bacterial communities from UV damage"

This file contains the following Supplementary Information:

- **Supplementary Analyses**
- **Supplementary Table S1-S4**
- **Supplementary Figures S1-S7**
- **Video S1**

### Supplementary Analysis

#### Pyoverdine expression rate of cell lineage types with varying degrees of UV damage

In the main text, we used time-lapse microscopy to study the impact of UV exposure on the cumulative expression of the pyoverdine synthesis gene *pvdA* in cell lineages varying in the degree of apparent UV-induced damage. We observed [A] healthy cells that started to divide and formed microcolonies; [B] moderately damaged cells that did not divide, but remained intact over the duration of the experiment; [C] severely damaged cells that started to divide, but eventually underwent explosive cell lysis, leading to the death of all cells in the microcolony; and [D] critically damaged cells that did not divide and eventually underwent cell lysis. Our analysis revealed that heavily damaged (lysing) cell lineages invested less into pyoverdine expression than less-damaged (non-lysing) cells. Given that the cumulative pyoverdine expression is calculated over the duration of the entire experiment, this analysis cannot exclude the possibility that increased expression levels of heavily damaged types are masked by their dying during the experiment. To account for this possibility, we repeated our analysis of pyoverdine expression, but focusing on the rate of pyoverdine expression, calculated by dividing the cumulative pyoverdine expression by the time over which it was detected.

We found that pyoverdine expression rates varied between cell lineage categories and media (Table S1B; Figure S6). However, this variation did not reflect increased rates of pyoverdine expression in heavily damaged as compared to healthy cell lineages, which featured comparable expression rates during their lifetime both in iron-limited and iron-rich medium (Table S3). Instead, we observed that critically damaged [D] types featured a lower expression rate than the less-damaged [C] types under iron limitation (Table S3). Moreover, both the severelydamaged [C] types and the critically damaged [D] types featured lower expression rates than the moderately damaged [B] types under iron-rich conditions (Table S3). Altogether, these results show that cells which apparently incurred substantial UV damage feature pyoverdine expression rates comparable to those of ‘healthy’ cells.

**Table S1 | Pyoverdine expression in UV exposed cell lineages.** Determinants of (A) cumulative pyoverdine expression and (B) the rate of pyoverdine expression by UV-exposed cell lineages.

| | **(A) cumulative pyoverdine expression** | | | **(B) pyoverdine expression rate** | | |
|---|---|---|---|---|---|---|
| **predictor** | $\chi^2$ | df | p | $\chi^2$ | df | p |
| Category | 589.01 | 3 | **< 0.001** | 23.94 | 3 | **< 0.001** |
| Medium | 98.11 | 1 | **< 0.001** | 85.42 | 1 | **< 0.001** |
| Category:Medium | 22.50 | 3 | **< 0.001** | 19.58 | 3 | **< 0.001** |

**Table S2 | Differences in cumulative pyoverdine expression among cell lineage types.** Post-hoc comparisons of cell lineage types varying in the degree of apparent UV-induced damage. P-values are adjusted for multiple testing using the false discovery rate. Significant p-values are in bold print.

| Medium | contrast | estimate | SE | df | t.ratio | p |
|---|---|---|---|---|---|---|
| iron-limited | healthy [A] - moderate [B] | 6.41 | 2.87 | 67.92 | 2.229 | **0.029** |
| iron-limited | healthy [A] - severe [C] | 16.21 | 3.47 | 73.69 | 4.671 | **< 0.001** |
| iron-limited | healthy [A] - critical [D] | 37.40 | 2.78 | 63.09 | 13.455 | **< 0.001** |
| iron-limited | moderate [B] - severe [C] | 9.81 | 3.30 | 98.57 | 2.971 | **0.005** |
| iron-limited | moderate [B] - critical [D] | 30.99 | 2.56 | 247.21 | 12.085 | **< 0.001** |
| iron-limited | severe [C] - critical [D] | 21.18 | 3.22 | 94.70 | 6.581 | **< 0.001** |
| iron-rich | healthy [A] - moderate [B] | -8.41 | 2.92 | 67.45 | -2.879 | **0.005** |
| iron-rich | healthy [A] - severe [C] | 20.77 | 3.12 | 80.11 | 6.655 | **< 0.001** |
| iron-rich | healthy [A] - critical [D] | 37.35 | 2.38 | 90.32 | 15.714 | **< 0.001** |
| iron-rich | moderate [B] - severe [C] | 29.18 | 3.45 | 88.75 | 8.461 | **< 0.001** |
| iron-rich | moderate [B] - critical [D] | 45.76 | 2.79 | 75.01 | 16.383 | **< 0.001** |
| iron-rich | severe [C] - critical [D] | 16.57 | 3.00 | 87.00 | 5.523 | **< 0.001** |

**Table S3 | Differences in the rate of pyoverdine expression among cell lineage types.** Post-hoc comparisons of cell lineage types varying in the degree of apparent UV-induced damage. P-values are adjusted for multiple testing using the false discovery rate. Significant p-values are in bold print.

| Medium | contrast | estimate | SE | df | t.ratio | p |
|---|---|---|---|---|---|---|
| iron-limited | healthy [A] - moderate [B] | 0.92 | 0.41 | 67.56 | 2.229 | 0.058 |
| iron-limited | healthy [A] - severe [C] | 0.07 | 0.49 | 74.96 | 0.138 | 0.891 |
| iron-limited | healthy [A] - critical [D] | 1.34 | 0.40 | 61.46 | 3.398 | **0.007** |
| iron-limited | moderate [B] - severe [C] | -0.85 | 0.46 | 108.01 | -1.844 | 0.102 |
| iron-limited | moderate [B] - critical [D] | 0.43 | 0.36 | 243.04 | 1.172 | 0.291 |
| iron-limited | severe [C] - critical [D] | 1.28 | 0.45 | 101.90 | 2.858 | **0.016** |
| iron-rich | healthy [A] - moderate [B] | -1.20 | 0.42 | 64.41 | -2.879 | **0.011** |
| iron-rich | healthy [A] - severe [C] | 0.51 | 0.40 | 84.71 | 1.274 | 0.247 |
| iron-rich | healthy [A] - critical [D] | 0.80 | 0.33 | 81.26 | 2.435 | **0.026** |
| iron-rich | moderate [B] - severe [C] | 1.71 | 0.45 | 79.74 | 3.797 | **< 0.001** |
| iron-rich | moderate [B] - critical [D] | 2.00 | 0.39 | 63.88 | 5143 | **< 0.001** |
| iron-rich | severe [C] - critical [D] | 0.29 | 0.37 | 102.87 | 0.785 | 0.434 |

**Table S4 | Structure of final models.** Models are given in the order in which they occur in the main text. Note that the response was square-root transformed in all models. The fixed effect structure was obtained by deleting non-significant interaction terms from the corresponding full model (including all possible interaction terms between explanatory variables). The appropriate variance structure was selected by fitting several models with all possible variance structures, and then selecting the model with the lowest AIC (see the Statistical Analyses section in the Materials and Methods of the main text for further details).

| **model** | **response [y]** | **fixed effect structure[1]** | **variance structure[2]** |
|---|---|---|---|
| main #1 | pyoverdine production | Supplement * Treatment * Medium | varIdent(form=~1\|Supplement*Medium) |
| main #2 | pyoverdine expression | Supplement + Treatment + Medium + Supplement:Treatment + Supplement:Medium | - |
| main #3 | growth | Strain * Supplement * Medium + Treatment + Supplement:Treatment | varIdent(form=~1\|Supplement*Treatment*Medium) |
| main #4 | duration of lag phase | Supplement * Treatment * Medium + Strain | varIdent(form=~1\|Supplement) |
| main #5 | mortality rate | Supplement * Treatment * Medium + Strain | varIdent(form=~1\|Supplement*Treatment*Medium) |
| main #6 | pyoverdine expression ['healthy' lineages] | Treatment + Medium | - |
| main #7 | pyoverdine expression [all lineages; after UV exposure] | Cell_lineage_category * Medium | varIdent(form=~1\|Cell_lineage_category*Medium) |
| supp #1 | pyoverdine expression rate [all lineages; after UV exposure] | Cell_lineage_category * Medium | varIdent(form=~1\| Cell_lineage_category*Medium) |

[1] A specification in the form of A+B indicates "main effect of A and main effect of B", whereas A:B indicates "interaction of A and B", and A*B indicates "main effect of A, main effect of B, and interaction of A and B"

[2] A specification in the form varIdent(form = ~1|A*B) indicates that the variance is allowed to differ for every combination of the different levels of A and B

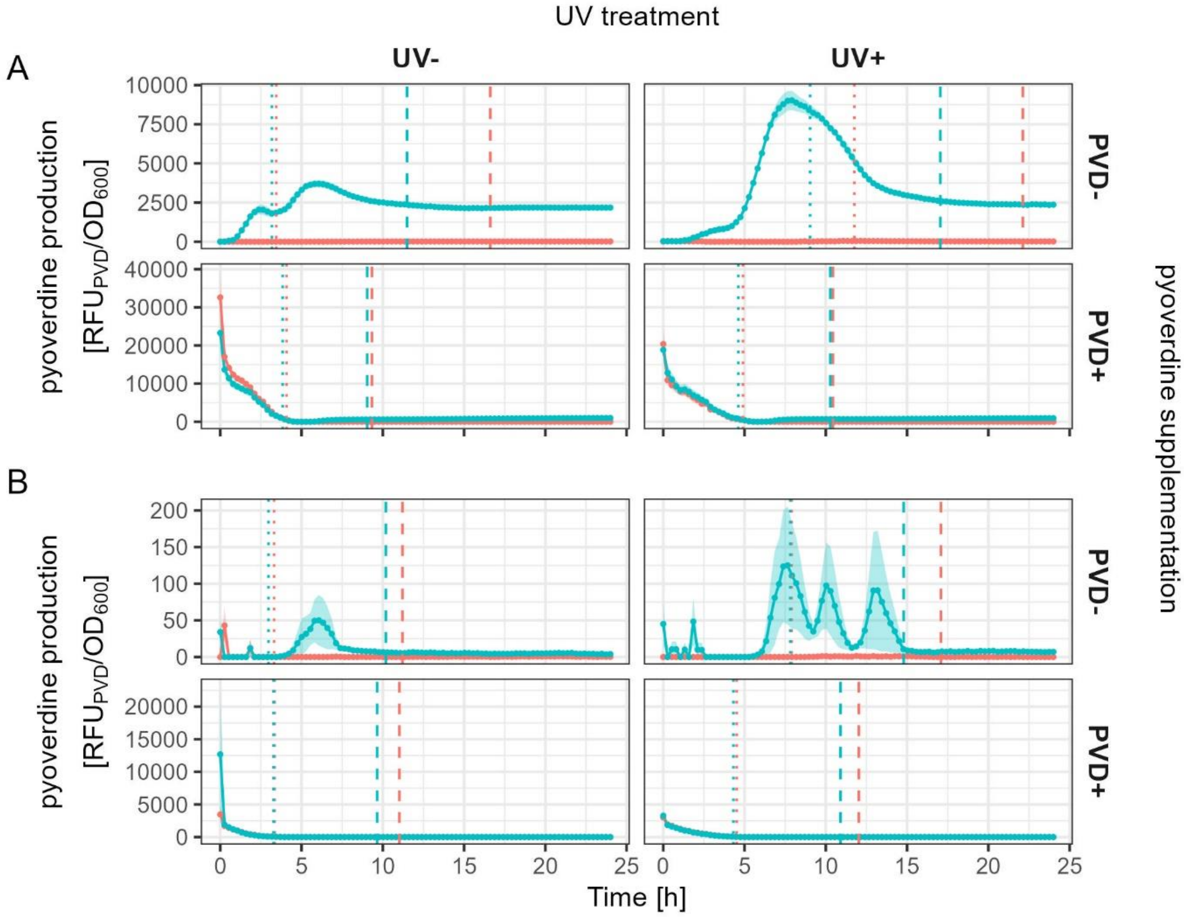


**Figure S1 | Per-cell production of pyoverdine over time.** Per-cell production of pyoverdine in monocultures of the *P. aeruginosa* PAO1 wildtype (turquoise) and the pyoverdine-deficient mutant PAO1Δ*pvdD* (red) grown under (A) iron-limited and (B) iron-rich conditions with or without prior UV exposure and with or without prior pyoverdine supplementation. Pyoverdine production was quantified as pyoverdine fluorescence [$RFU_{PVD}$] divided by growth [$OD_{600}$]. Dotted and dashed vertical lines denote the end of the lag-phase and the end of the exponential growth phase (determined by for each strain based on its growth; see Figure S3), respectively. Solid lines depict the mean across replicates, colored bands show the standard error. Note that the y-axis range differs between panels to properly resolve variation between strains across treatment combinations.

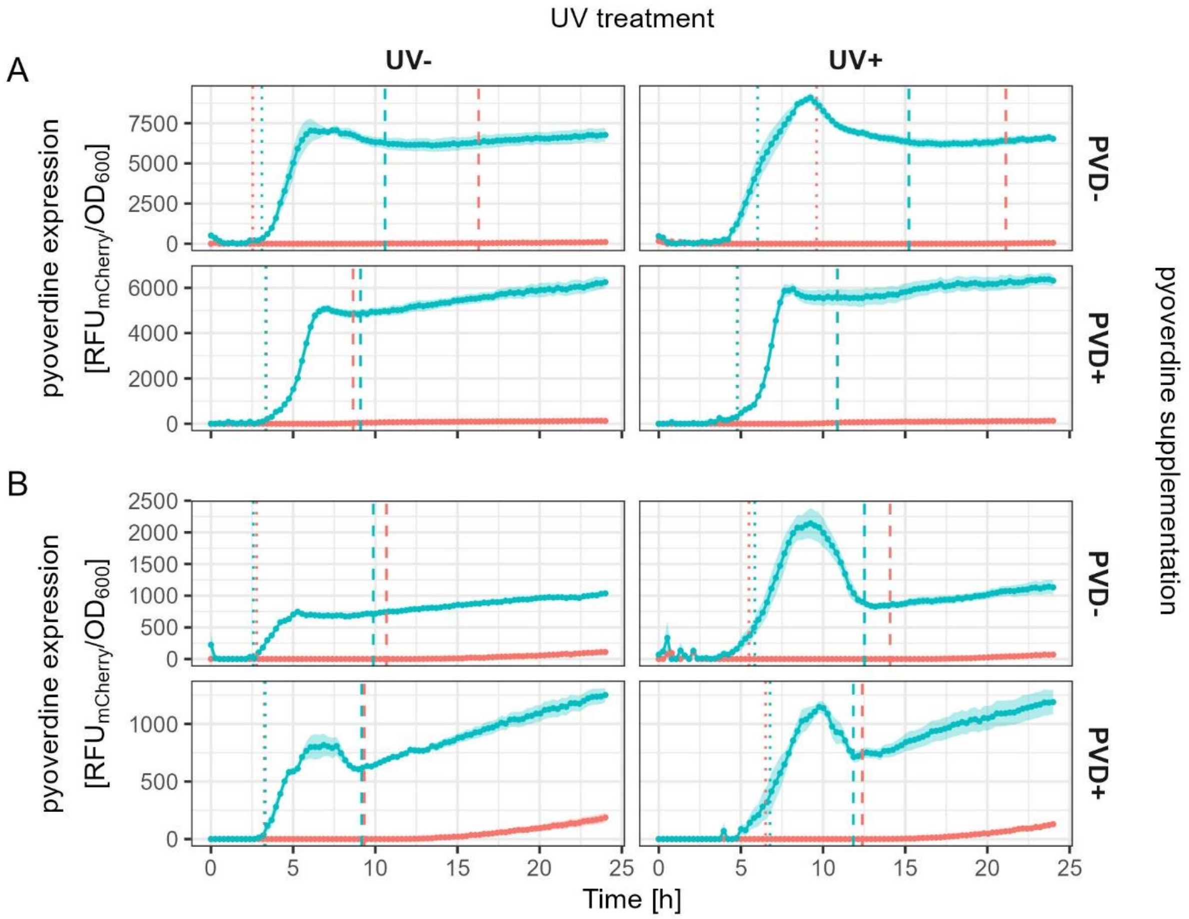


**Figure S2 | Per-cell expression of the pyoverdine synthesis gene *pvdA* over time.** Per-cell expression of pyoverdine in monocultures of the *P. aeruginosa* reporter strain PAO1*pvdA::mCherry* (turquoise) and the pyoverdine-deficient mutant PAO1Δ*pvdD* (red) grown under (A) iron-limited and (B) iron-rich conditions with or without prior UV exposure and with or without prior pyoverdine supplementation. Pyoverdine expression was quantified as mCherry fluorescence [$RFU_{mCherry}$] divided by growth [$OD_{600}$]. Dotted and dashed vertical lines denote the end of the lag-phase and the end of the exponential growth phase (determined by for each strain based on its growth; see Figure S3), respectively. Solid lines depict the mean across replicates, colored bands show the standard error. Note that the y-axis range differs between panels to resolve variation between strains across treatment combinations.

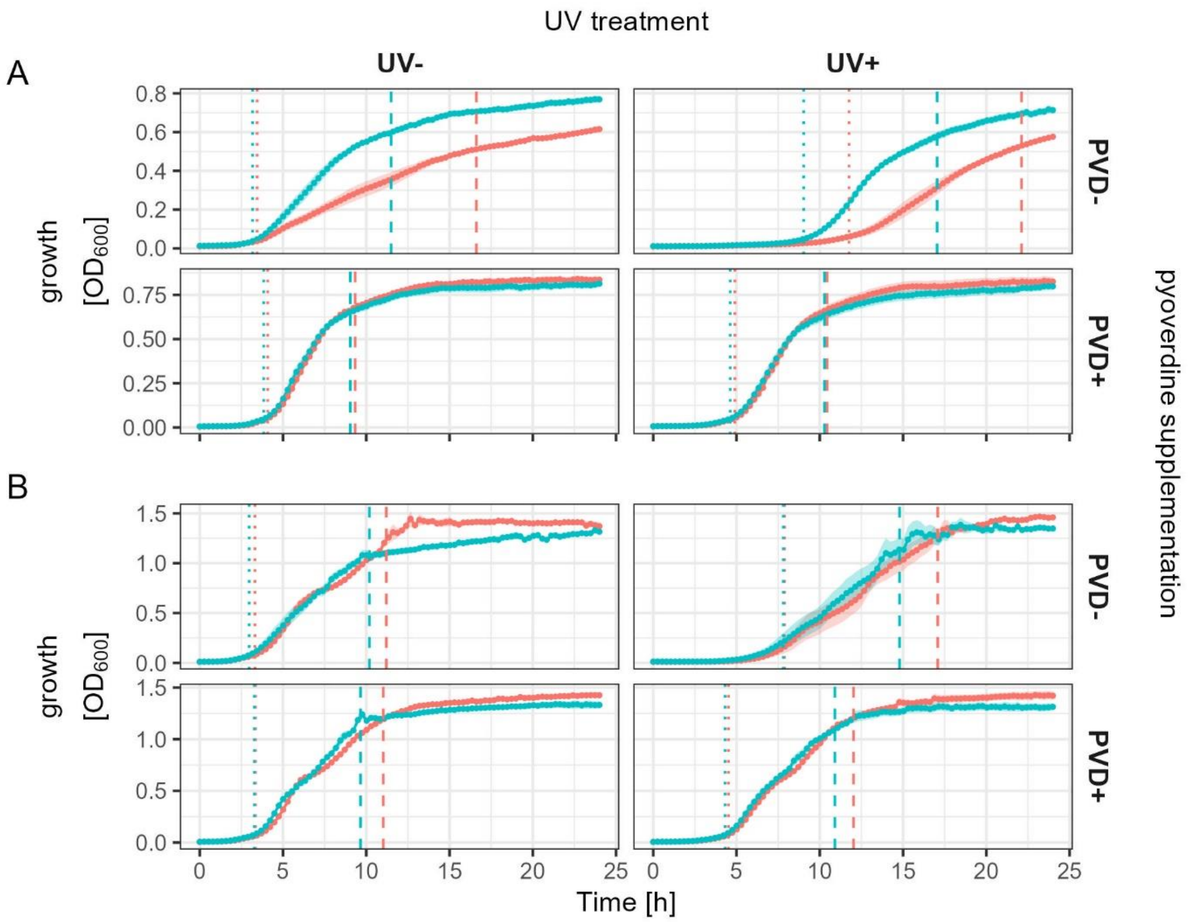


**Figure S3 | Growth over time.** Monoculture growth of the *P. aeruginosa* PAO1 wildtype (turquoise) and the pyoverdine-deficient mutant PAO1Δ*pvdD* (red) in (A) iron-limited and (B) iron-rich conditions with or without prior UV exposure and with or without prior pyoverdine supplementation. Growth was quantified as optical density [$OD_{600}$]. Dotted and dashed vertical lines denote the end of the lag-phase and the end of the exponential growth phase, respectively. The indicated durations of growth phases represent the mean of values extracted from Gompertz-models fit on individual replicates. Solid lines depict the mean across replicates, colored bands show the standard error. Note that the y-axis range differs between panels to properly resolve variation between strains across treatment combinations.

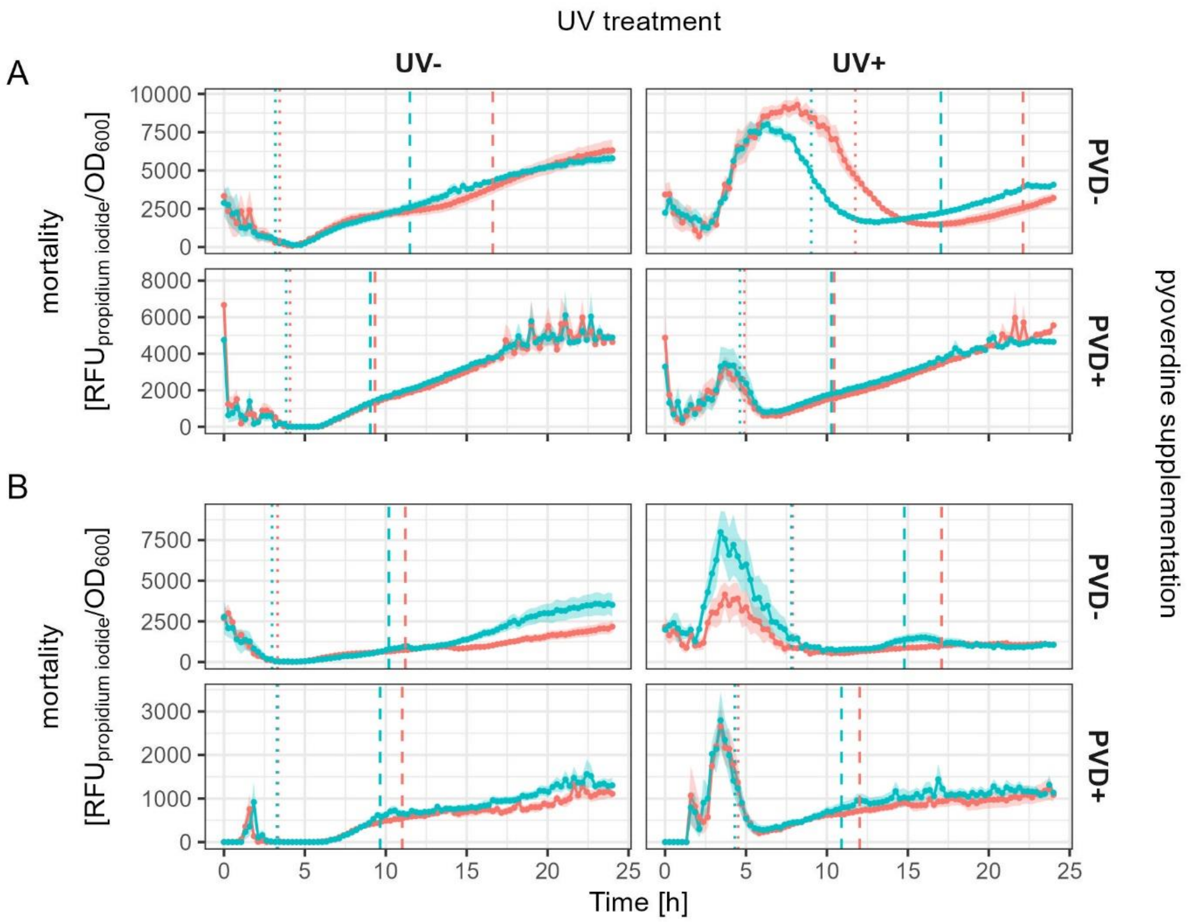


**Figure S4 | Mortality over time.** Mortality in monocultures of the *P. aeruginosa* PAO1 wildtype (turquoise) and the pyoverdine-deficient mutant PAO1Δ*pvdD* (red) grown under (A) iron-limited and (B) iron-rich conditions with or without prior UV exposure and with or without prior pyoverdine supplementation. Mortality was quantified as propidium iodide fluorescence [$RFU_{propidium\ iodide}$] divided by growth [$OD_{600}$]. Dotted and dashed vertical lines denote the end of the lag-phase and the end of the exponential growth phase (determined by for each strain based on its growth; see Figure S3), respectively. Solid lines depict the mean across replicates, colored bands show the standard error. Note that the y-axis range differs between panels to properly resolve variation between strains across treatment combinations.

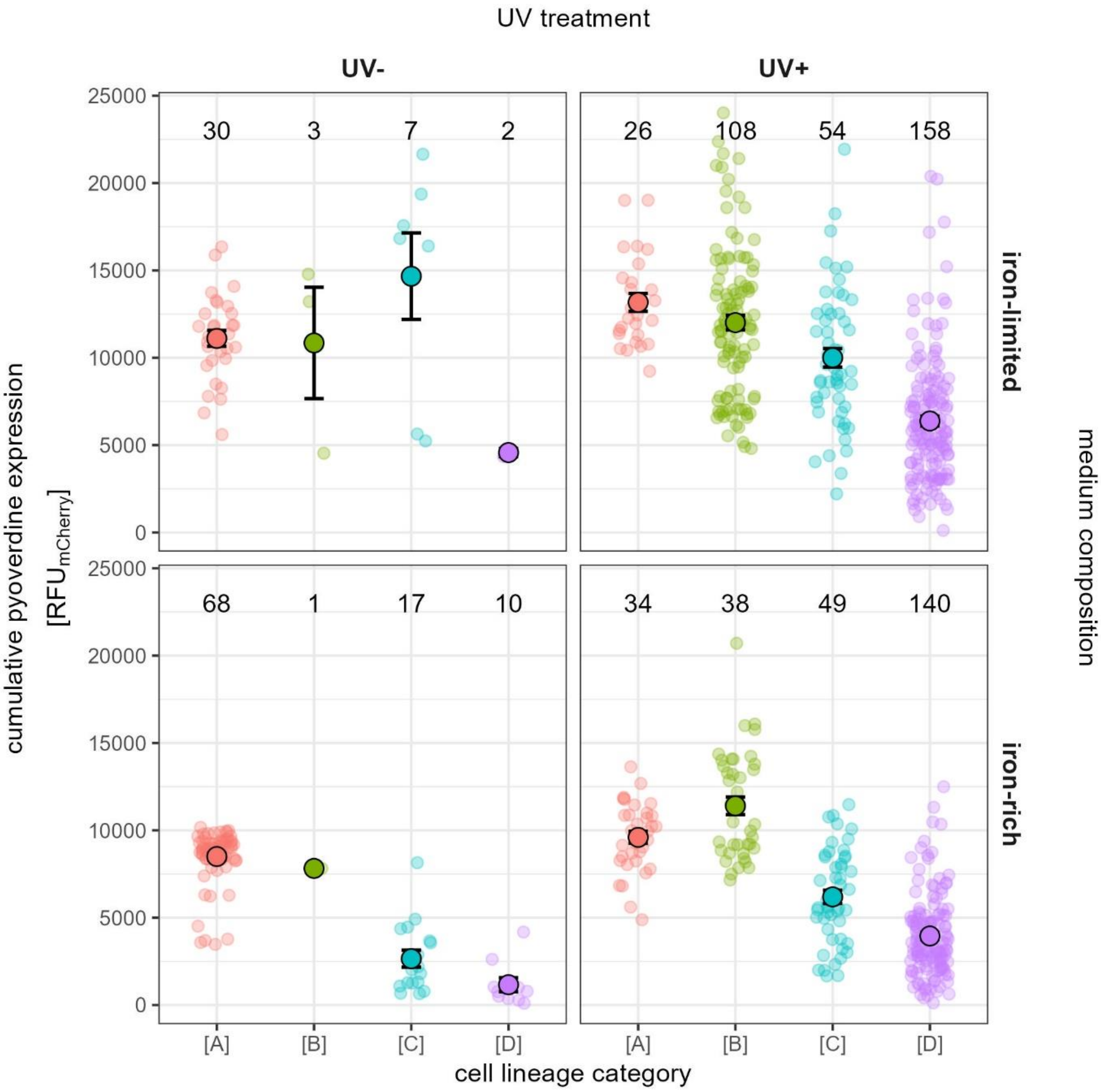


**Figure S5 | Cumulative expression of the pyoverdine synthesis gene *pvdA* in different cell lineage types.** Cumulative expression of the pyoverdine synthesis gene *pvdA* in different cell lineage types varying in the degree of apparent UV damage, measured via single-cell microscopy under iron-limited and iron-rich conditions with or without prior UV exposure. Pyoverdine expression was quantified as the integral of mCherry fluorescence measured over the whole duration of the experiment (8h). Cell lineage types vary in the degree of apparent UV-induced damage: [A] healthy cells (red) start to divide and form microcolonies; [B] moderately damaged cells (green) do not divide, but remain intact over the duration of the experiment; [C] severely damaged cells (blue) start to divide, but eventually undergo explosive cell lysis, leading to the death of all cells in the microcolony; [D] critically damaged cells (lilac) do not divide and eventually undergo explosive cell lysis. Small circles are individual replicates, and either represent the signal of individual cells (for non-dividing types [B] and [D] featuring moderate or critical damage) or the average signal of all cells belonging to the same lineage (for dividing types [A] and [C] featuring no or severe damage). Large circles and black lines show mean and standard error. Numbers within panels indicate sample sizes. Note that we did not statistically analyze the pyoverdine expression levels of damaged phenotypes in the absence of prior UV exposure due to their very low frequency.

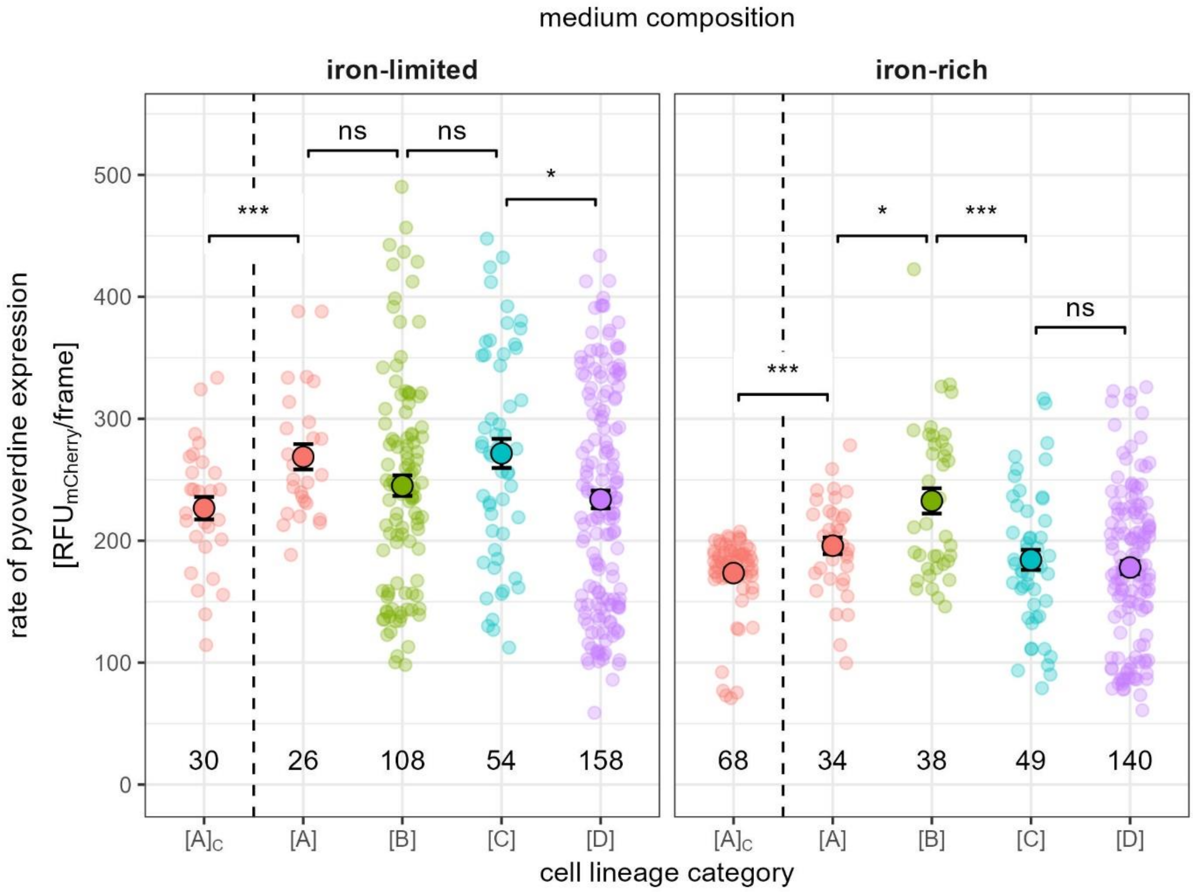


**Figure S6 | Expression rate of the pyoverdine synthesis gene *pvdA* after UV exposure in different cell lineage types.** The rate of pyoverdine expression was quantified under iron-rich and iron-limited conditions as the integral of mCherry fluorescence divided by the number of frames that the focal cell (lineage) survived. Cell lineage types vary in the degree of apparent UV-induced damage: [A] healthy cells (red) start to divide and form microcolonies; [B] moderately damaged cells (green) do not divide, but remain intact over the duration of the experiment; [C] severely damaged cells (blue) start to divide, but eventually undergo explosive cell lysis, leading to the death of all cells in the microcolony; [D] critically damaged cells (lilac) do not divide and eventually undergo explosive cell lysis. $[A]_C$ are healthy cells (red) from cultures without UV exposure and serve as a negative control. Small circles are individual replicates, and either represent the signal of individual cells (for non-dividing types [B] and [D] featuring moderate or critical damage) or the average signal of all cells belonging to the same lineage (for dividing types [A] and [C] featuring no or severe damage). Large circles and black lines show mean and standard error. Numbers within panels indicate sample sizes. Significance levels of post-hoc comparisons are indicated as follows: * $0.05 \geq p > 0.01$; ** $0.01 \geq p > 0.001$; *** $p \leq 0.001$.

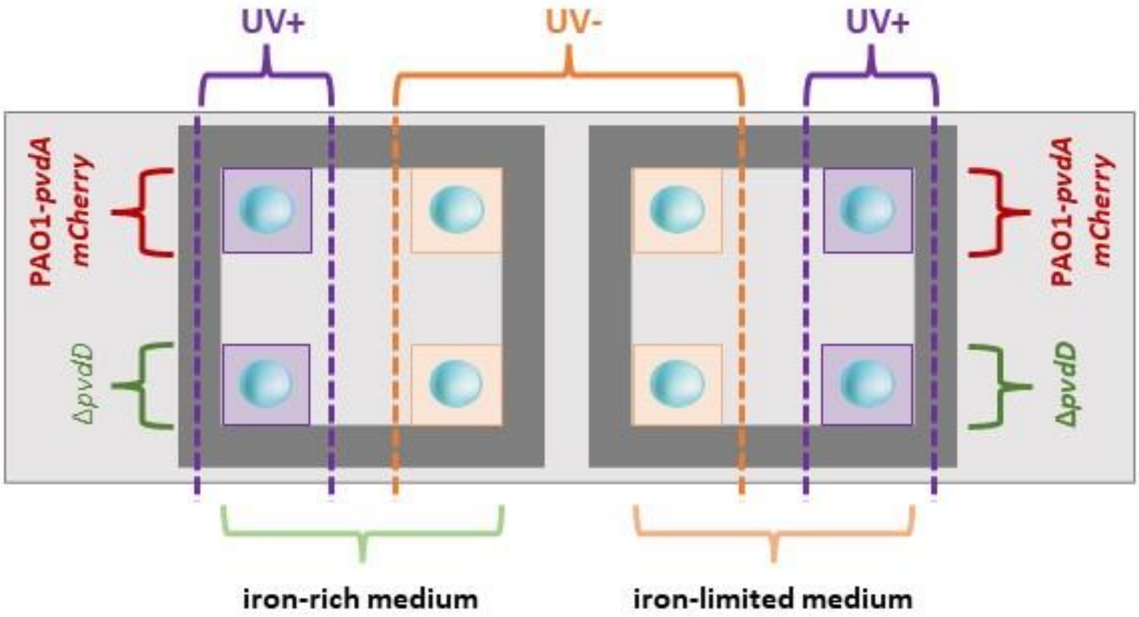


**Figure S7 | Microscopy slide design.** Design of the microscopy slide setup used to quantify heterogeneity in pyoverdine expression at the single-cell level. Please note that the pyoverdine-deficient mutant PAO1 Δ*pvdD* was only used as an internal control.

**Video S1 | Cell lineage types after UV exposure.** The accompanying video file ("VideoS1_CellLineageTypes") shows four cell lineage types varying in the degree of apparent UV-induced damage: [A] healthy cells start to divide and form microcolonies; [B] moderately damaged cells do not divide, but remain intact over the duration of the experiment; [C] severely damaged cells start to divide, but eventually undergo explosive cell lysis, leading to the death of all cells in the microcolony; and [D] critically damaged cells do not divide and eventually undergo explosive cell lysis.
